# A wipe-based stool collection and preservation kit for microbiome community profiling

**DOI:** 10.1101/2021.12.03.471072

**Authors:** Hui Hua, Cem Meydan, Evan E. Afshin, Loukia Lili, Christopher R. D’Adamo, Joel Dudley, Nathan D. Price, Bodi Zhang, Christopher E. Mason

## Abstract

While a range of methods for stool collection exist, many require complicated, self-directed protocols and stool transfer. In this study, we introduce and validate a novel, wipe-based approach to fecal sample collection and stabilization for metagenomics analysis. A total of 72 samples were collected across four different preservation types: freezing at -20°C, room temperature storage, a commercial DNA preservation kit, and DESS (dimethyl sulfoxide, ethylenediaminetetraacetic acid, sodium chloride) solution. These samples were sequenced and analyzed for taxonomic abundance metrics, metabolic pathway classification, and diversity analysis. Overall, the DESS wipe results validated the use of a wipe-based capture method to collect stool samples for microbiome analysis, showing an R^2^ of 0.96 for species across all kingdoms, as well as exhibiting a maintenance of Shannon diversity (3.1-3.3) and species richness (151-159) compared to frozen samples. Moreover, DESS showed comparable performance to the commercially available preservation kit (R^2^ of 0.98), and samples consistently clustered by subject across each method. Future studies will be needed to further explore sample processing options and their applications in non-healthy subjects, particularly patients with irritable bowel syndrome, inflammatory bowel disease, and colorectal cancer, but these data suggest the DESS wipe method can be used for stable, room temperature collection and transport of human stool specimens.

## Introduction

The detection and identification of biomolecules in microbial communities from samples is widely used for monitoring disease and overall health. Stable transportation and delivery of biomolecules is generally required for such analysis. As such, low cost and efficient collection, storage, and delivery of biomolecules are critical for the field of medical diagnosis.

For human gut microbiome analysis, recent advances in sequencing techniques and bioinformatics have increased our knowledge of the complex microbial communities and their interactions. It is now well established that these microbes play important roles in relation to inflammation [1], metabolic disease [2, 3], mental disorders [4, 5], aging [6, 7] and several other diseases and health conditions [8-12]. However, different approaches of sample processing can introduce human error variability or technical biases through inappropriate sample handling or storage. For example, fecal microbiota sequencing profiles have been shown to change significantly during ambient temperature storage after 48 hours [13,14]. While performing nucleic acid extraction on fresh samples immediately after collection is impractical, freezing and storing samples at −20 °C, −80 °C, or below, is widely considered to be best practice when preserving microbial composition for sequence-based analysis [15–17]. However, this is difficult to achieve in many situations, such as sampling in remote areas, and thus may dramatically increase the costs of such studies. While some studies have investigated in detail the rapid deterioration of fecal samples that have been stored at room temperature for several days prior to lab processing [13, 18–20], there are few methods to address such issues.

Moreover, stool specimen collection using most methods can be challenging, and many individuals find the process difficult and not user-friendly [21]. Challenges include embarrassment, fear of results, concerns around hygiene and contamination, discretion and privacy, and lack of information. A 3-year randomized trial of 997 participants found that discomfort with the collection of a stool sample is the most frequently cited barrier for participation in fecal test-based screening. Furthermore, the study found that having a choice of screening methods significantly increases (13% vs. 43%) patient adherence [22].

A 2016 study, spanning 15 individuals and over 1,200 samples, provided the most comprehensive view to date of storage and stabilization effects on stool [23]. It suggested that 95% ethanol can preserve samples sufficiently well at ambient temperatures for periods of up to 8 weeks, and include the types of variation often encountered under field conditions, such as freeze-thaw cycles and high temperature fluctuations [23]. In addition, a solution containing dimethyl sulphoxide, disodium EDTA, and saturated NaCl (DESS) was originally used for various applications in the preservation of nematodes for combined morphological and molecular analyses, has also been used to preserve entire soil/sediment samples, or as a storage medium for microbial community analysis [24-26]. Such preserved material can be easily stored for months at room temperature, shipped by mail, or carried in luggage, which provides an efficient, cost-effective method with widespread applications for microbiome studies.

To address such technical errors and biases in sample collection methods, as well as to enhance the user experience of stool sample collection, we have designed a practical and user-friendly fecal sample collection kit that includes a dissolvable wipe (e.g., ethanol-soluble film), in which the biological sample is dissolved in a DNA stabilizing solution (DESS). The film, solution, and biological sample are disposed of in a sealable container. Therefore, people can collect their fecal samples as easily as using toilet paper after defecation. The DNA stabilizing solution ensures that the microbiomic community structure is well-preserved during ambient temperature transportation and storage, and the microbiome DNA is extracted from the fixed microbes and used for further laboratory analysis when the samples arrive at the laboratory. In addition, the dissolvable characters of the wipe ensure that all the microbes contact the stabilizing solution adequately when the sample is collected. The primary objective of this study is to assess the extent to which our novel approach to fecal sample collection and stabilization could maintain microbiota composition relative to immediate freezing (−20°C), preservation with a commercially available kit, and storage at room temperature (RT).

## Results

### Study Summary

Figure 1. summarizes the overall study design. Six subjects were recruited to participate in the study to validate the wipe capture method. Two males and four females enrolled in the study. The median age was 42 years old (range: 31-60 years). Three subjects were white, non-Hispanic and three subjects were black, non-Hispanic. The average BMI across the cohort was 27.8 kg/m^2^ (range: 20.3-40.7 kg/m^2^). Four preservation methods were used to process the samples for metagenomics sequencing: freezing (−20°C), room temperature storage, a commercial preservation kit (room temperature), and DESS DNA preservation (room temperature). Three replicates per subject for each preservative were collected for a total of 72 samples. A total of 71 samples were successfully sequenced with an average of 8 million sequencing reads and a range of 3 to 38 million reads. One sample was removed from the analysis due to sequencing failure (no library). The amount of DNA captured by wipe was found to be comparable to other collection and extraction methods. The average DNA yields from extraction were 98.2ng/uL for DESS, 44.6ng/uL for the commercial preservation kit, 286ng/uL for the -20°C samples, and 122.3ng/uL for the room temperature samples.

**Figure 1.**
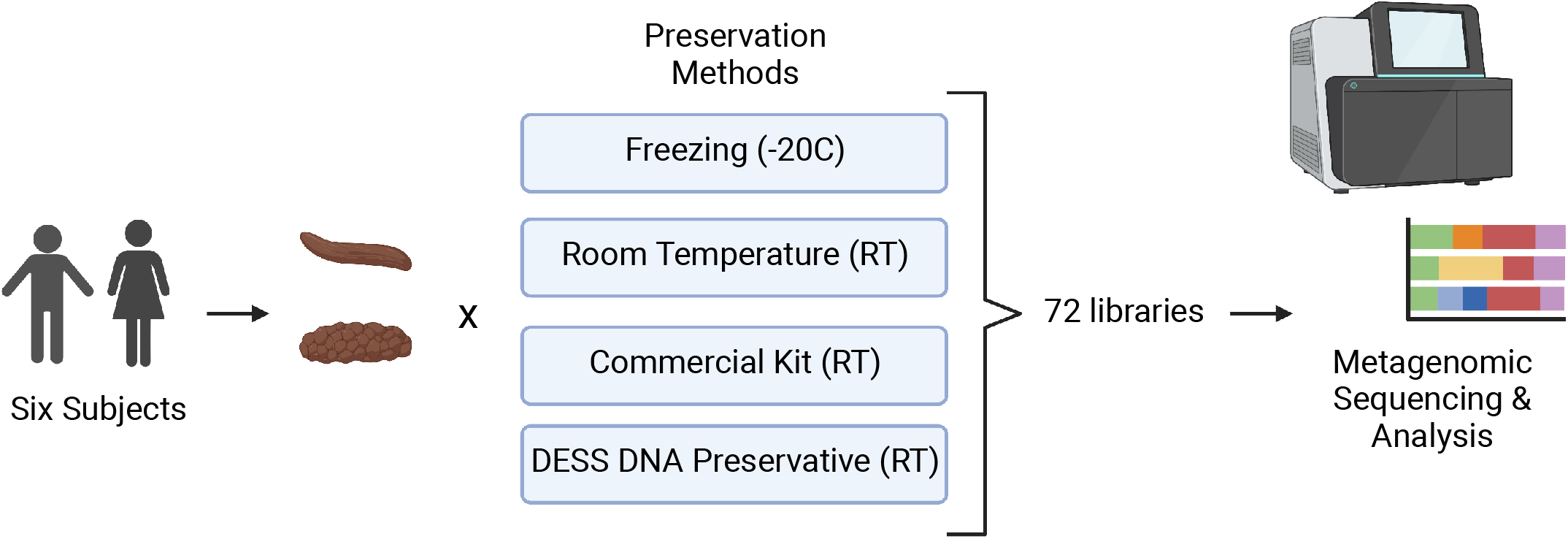
Study Design. Six participants were enrolled in the study and collected stool samples (Bristol Scale Type 3 and 4) for metagenomics/microbiome analysis. The samples were processed using four different preservation techniques: freezing at -20°C, stored at room temperature (RT). Zymo DNA Shield (RT), and DESS (RT). A total of 72 samples were then sequenced with next-generation sequencing and analyzed for taxa and metabolic profiles. Created with BioRender.com.

### Sample Similarity

A t-SNE analysis across sample types and subjects showed clear clustering by each individual across the cohort, wherein each subject was isolated and separated from one another based on their unique microbiome signature (**Figure 2**). Interestingly, wipe samples in the DESS clustered more closely with the frozen samples and the commercial preservation kit’s samples. Meanwhile, in most subjects, the negative control samples that were stored at room temperature cluster together separately from the other preservation types. **Supplemental Figure 1** further demonstrates this finding in a dendrogram showing clustering by subject and divergence of room temperature samples compared to the other sample types.

**Figure 2.**
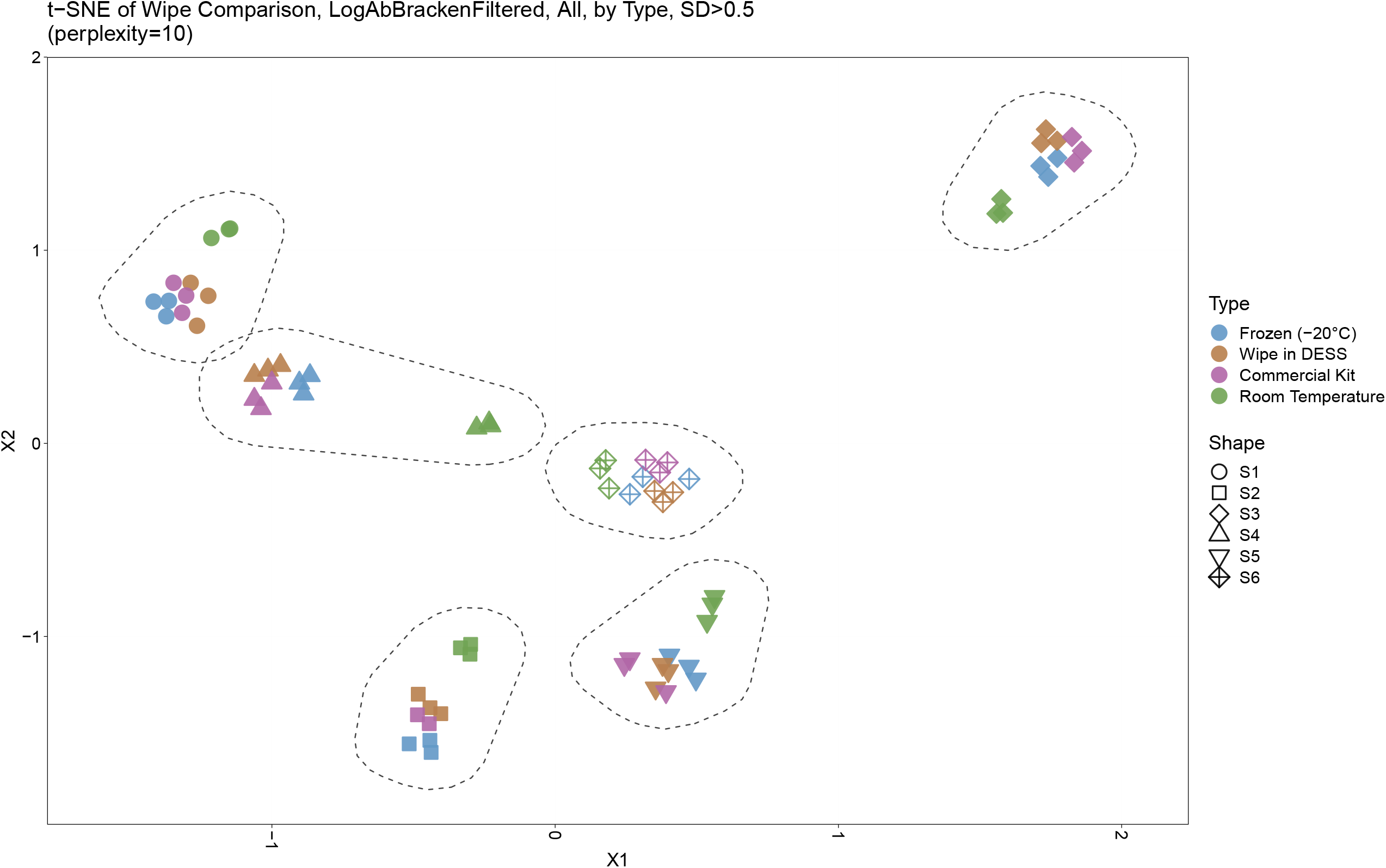
Sample Similarity. A t-SNE plot displaying sample comparisons and clustering. Sample types are denoted by different colors and subjects by different shapes. Six distinct clusters are shown, one for each subject, and froze, wipe in DESS, and commercially processed samples cluster together while room temperature samples cluster separately from the other preservation types.

### Taxonomic Profiles

Taxonomic assignment of reads to each domain of life were then examined for their relative distributions across the sample types. As expected, Bacteria was the predominant domain (>99% relative abundance) captured by the microbiome analysis across all subjects (**Figure 3A**). Subject 1 had some more hits to Archaea (<l%) than others, Subjects 4 and 6 had some samples with Eukaryota hits, and Subject 5 had some samples with viral hits (most <1%) (**Figure 3B)**. Commensal gut flora including *Firmicutes, Bacteroidetes*, and *Actinobacteria*, were the top phyla across all samples, with some room temperature samples also having *Proteobacteria*, particularly in Subject 4. **Figures 3C** and **3D** highlight the correlation of the subject’s microbiome profile across different domains comparing wipe DESS preparation to frozen and the commercial kit to frozen, respectively. There is an increased relative abundance of human DNA seen in wipe samples compared to the commercial kit (**Figure 3C**), however, this is still negligible compared to the predominance of reads matching to bacteria (**Figure 3A**).

**Figure 3.**
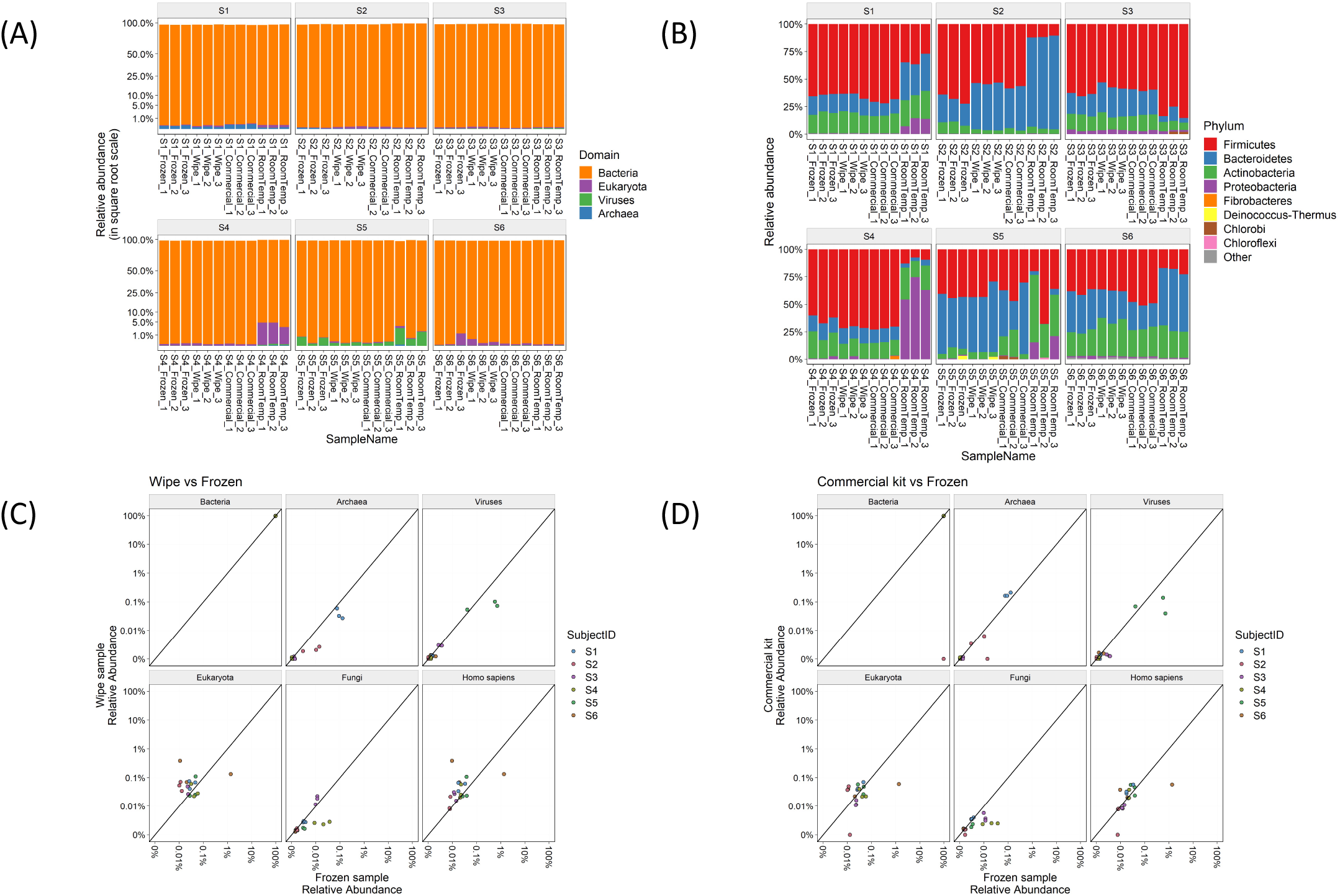
Taxonomic Profiles. Relative abundances of (A) Domains and (B) Phyla across the different subjects and sample types. Correlation plots comparing the relative abundances of wipe in DESS vs frozen samples (C) and commercial DNA preservation vs frozen samples (D).

### Diversity Metrics

The metagenomic data were then examined for two metrics of species diversity (**Figure 4**). The Shannon index metric showed a similar range (2.7-3.8) across all sample types, but the wipe in DESS (median 3.2) had more comparable levels to the commercial preservation method (3.1) and gold standard frozen samples (3.2), than the room temperature storage (2.9). The median species richness (151-159), however, was more comparable across all preservation techniques (**Figure 4B**).

**Figure 4.**
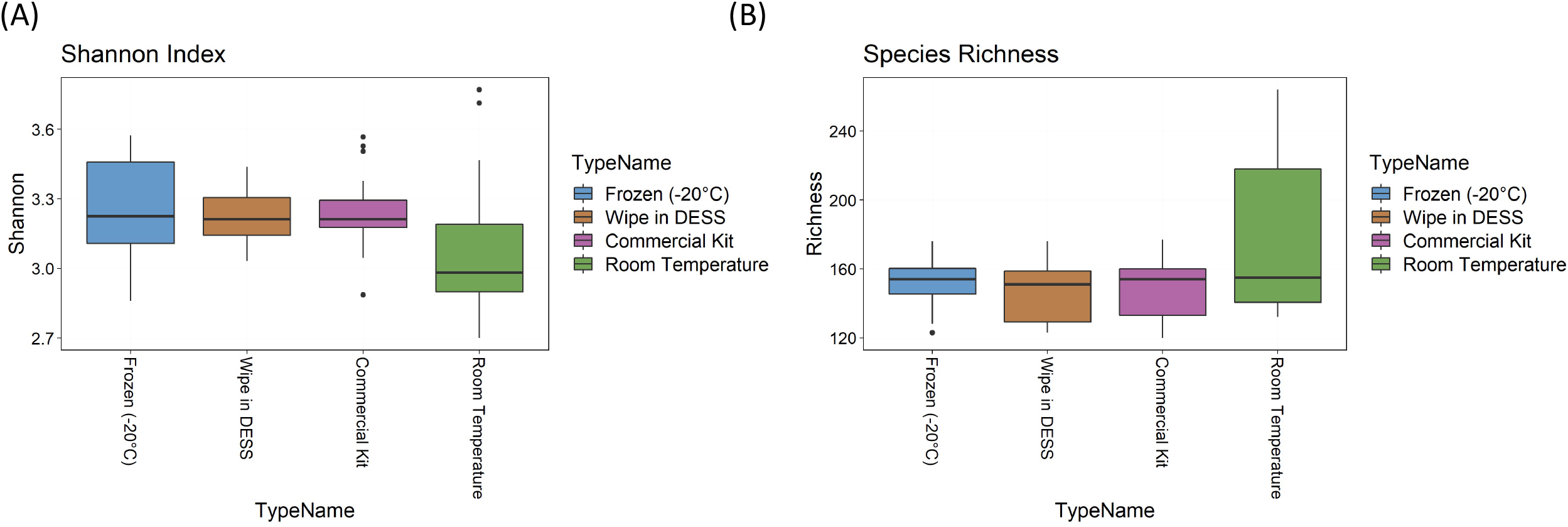
Diversity Metrics. (A) Shannon index and (B) Species richness diversity metrics across sample types.

### Intra- and Inter-sample Comparisons

Taxonomic profiles comparing the different preservation methods showed that DESS has a very strong correlation with the frozen samples and is comparable to the commercial kit (**Figure 5**). The Pearson correlation of taxa log abundance with intra-group and inter-group comparisons. DESS was found to be very similar to the -20°C frozen samples when considering replicate-to-replicate variability (positive-to-positive correlation = 0.92, positive-to-DESS correlation = 0.91) (**Figure 5A**). Furthermore, **Figures 5B** and **5C** highlight Pearson correlations calculated by median log10 relative abundances and median HUMAnN functional pathway scores, respectively (**Supplemental Figures 2** and **3**). These analyses demonstrate strong intra-sample correlation across the wipe samples, as well as strong inter-sample correlation between the wipe and frozen samples. These correlations are even comparable to the correlation found between the commercial preservation and frozen samples (**Figures 5B, 5C**). **Figures 5D** and **5E** highlight each subject’s taxonomic relative abundance comparing wipe to frozen and commercial preservation to frozen samples respectively. They further demonstrate that the wipe in DESS preservation method has a high positive correlation with the taxonomic profiles of the gold standard frozen preservation, and is comparable to the commercial preservation method.

**Figure 5.**
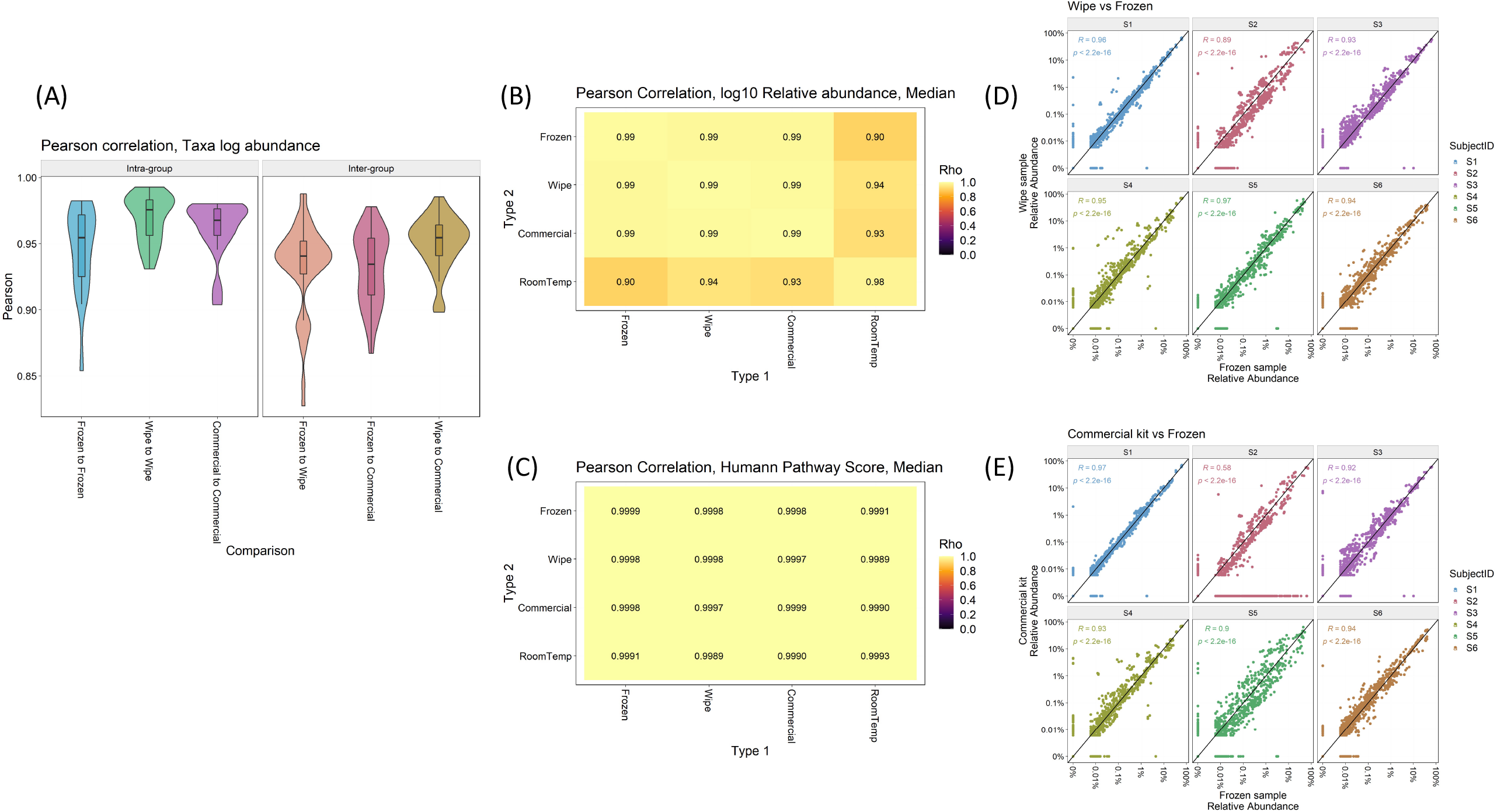
Intra- and Inter-sample Sample Comparisons. Pearson correlation by (A) Taxa log abundance with intra- and inter-group comparisons, (B) Median log10 relative abundances, (C) Median HUMAnN pathway scores, and correlation plots comparing the relative abundances found in (D) Wipe vs Frozen and (E) Commercial preservation vs Frozen samples.

## Discussion

This study demonstrates the use of a wipe-based capture method to collect stool samples for microbiome/metagenomics analysis. The DESS wipe preservation method showed comparable performance to a commercial DNA/RNA preservation kit, and was also very similar (R^2^ > 0.96) to the gold standard frozen samples for most metrics (i.e. taxonomic classification, diversity, functional pathway classification, and abundances). Both the DESS and the commercial preservation method showed significant diversity compared to the room temperature negative control.

Although the quality and significance of standard microbiome metrics are comparable across the wipe method, gold standard, and other commercial methods, further validation and better understanding of the bacterial to human DNA ratio in a broader population can be addressed in future studies. This will involve including non-healthy subjects such as samples from people with gut conditions (i.e. bloody diarrhea, IBS, blood in the stool, colorectal cancer, hemorrhoids, etc.) where human DNA is more present in the stool [27, 28], to further assess the performance of the wipe and integrity of microbiome analysis. Recruiting subjects with GI conditions such as constipation and diarrhea will further test the efficacy of the wipe. Moreover, subjects with different disease statuses and infections will be important to test, specifically patients with irritable bowel syndrome, IBD, *Clostridium difficile* infection, etc. Finally, RNA preservation and isolation for metatranscriptomics analysis poses its own set of unique challenges [29] and future studies will be needed to assess the wipe capture in DESS preservation for RNAseq analysis.

However, this study shows evidence of validation for a wipe-based collection and RT transport method for gut microbiome sampling and metagenomics sequencing analysis. Such a method may enable easier access to sampling, testing, and metagenomics implementation in clinical trials, home use, or even in remote environments, especially given the stability of the method. Indeed, wipe-based collection and processing offers a more user-friendly approach to collecting stool samples for microbiome analysis. Its ease-of-use design and simple instructions (just wipe and place into the tube) should enable easy integration with commercial stool collection kits and future biomedical studies and trials. Indeed, tools and methods such as these can be applied to help deploy metagenomics tools and methods for a wide range of both research and clinical applications.

## Methods

### Study Design

A total of 6 subjects were enrolled in this pilot study. The inclusion criteria to be enrolled in the study included: age ≥18; Bristol Stool Scale type 3 and 4 (normal), agree to collect and donate the feces, and the ability to understand and write English. Exclusion criteria included people with constipation, slightly dry, or diarrhea feces (Bristol Stool Scale types 1-2, 5-7), pregnant or breastfeeding females, history of alcohol, drug, or medication abuse, known allergies to any substance in the study product, current diagnosis of inflammatory bowel disease (Crohn’s Disease or Ulcerative Colitis), and currently taking any medication that may interfere with defecation.

### Sample Collection and Processing

Fecal collection kits were created and mailed to enrolled subjects with clear instructions on sample collection. A total of 12 samples were collected by each subject yielding a total of 72 samples to be processed. Four preservation methods were used to process the samples for metagenomics sequencing: freezing (−20°C), room temperature storage, Zymo DNA shield kit (room temperature), and DESS DNA preservation (room temperature). All the samples are shipped at room temperature except the samples meant for freezing which were shipped on dry ice. It took several days to up to a week for the shipment. When the samples arrived in the lab, all the samples were put into a fridge (4°C) except the frozen samples which were put in a -20°C freezer.

### Microbiome Sequencing and Analysis

DNA was extracted from all samples using QIAgen PowerSoil Pro Kit, libraries were prepared with Illumina Nextera FLEX, and samples were sequenced on the NextSeq500 platform. Samples were sequenced as paired-end 150bp for a mean depth of 8.0 million reads per sample (min: 2.9M, max:38.9M). Resulting sequences were trimmed by Trimmomatic [30], and then aligned to human genome reference using bwa [31]. Taxonomic annotation was performed by utilizing KrakenUniq [32] and subsequently Bracken [33] on a database that includes all bacterial, archaeal, viral, fungal references from RefSeq along with human reference. The lowest common ancestor taxonomic annotations were adjusted within the lineage until at least 10% of the unique k-mers belong to a specific clade and not its parent, then filtered for at least 10 reads and a minimum Bracken adjusted relative abundance of 0.005%. The pathway annotations were performed by using HUMAnN3 with the UniRef90 clusters [34].

## IRB

The study protocol was approved by the Institutional Review Board of the University of Maryland Baltimore (Protocol # HP-00087571). The study participants provided their written informed consent before enrolling in the study.

## Author Contributions

HH, CM, BZ, JTD, and CEM designed the study. HH, BZ, and CRD coordinated the IRB and subject recruitment. HH, CM, EA, LL, NDP, BZ, CEM were involved in data analysis, interpretation, and preparation of the manuscript. All authors approved and contributed to writing the manuscript.

## Competing Interests

All authors except for CRD are employees of Thorne HealthTech and the Onegevity Health platform.

