## Supplemental Materials for "A wipe-based stool collection and preservation kit for microbiome community profiling"

**Supplemental Figures**

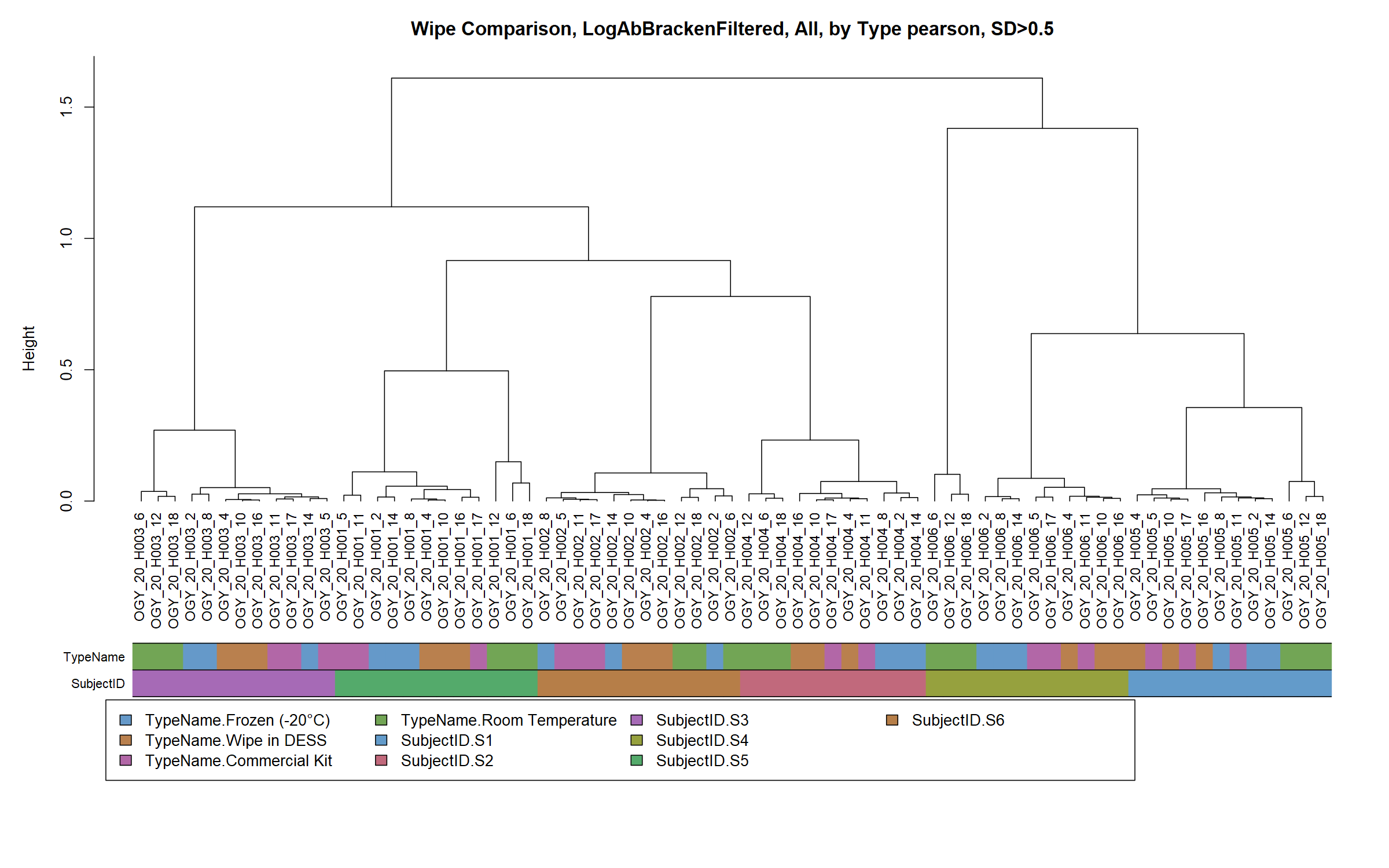

**Supplemental Figure 1. Sample Clustering.** Dendrogram plot showing sample clustering and similarity by sample type and subject.

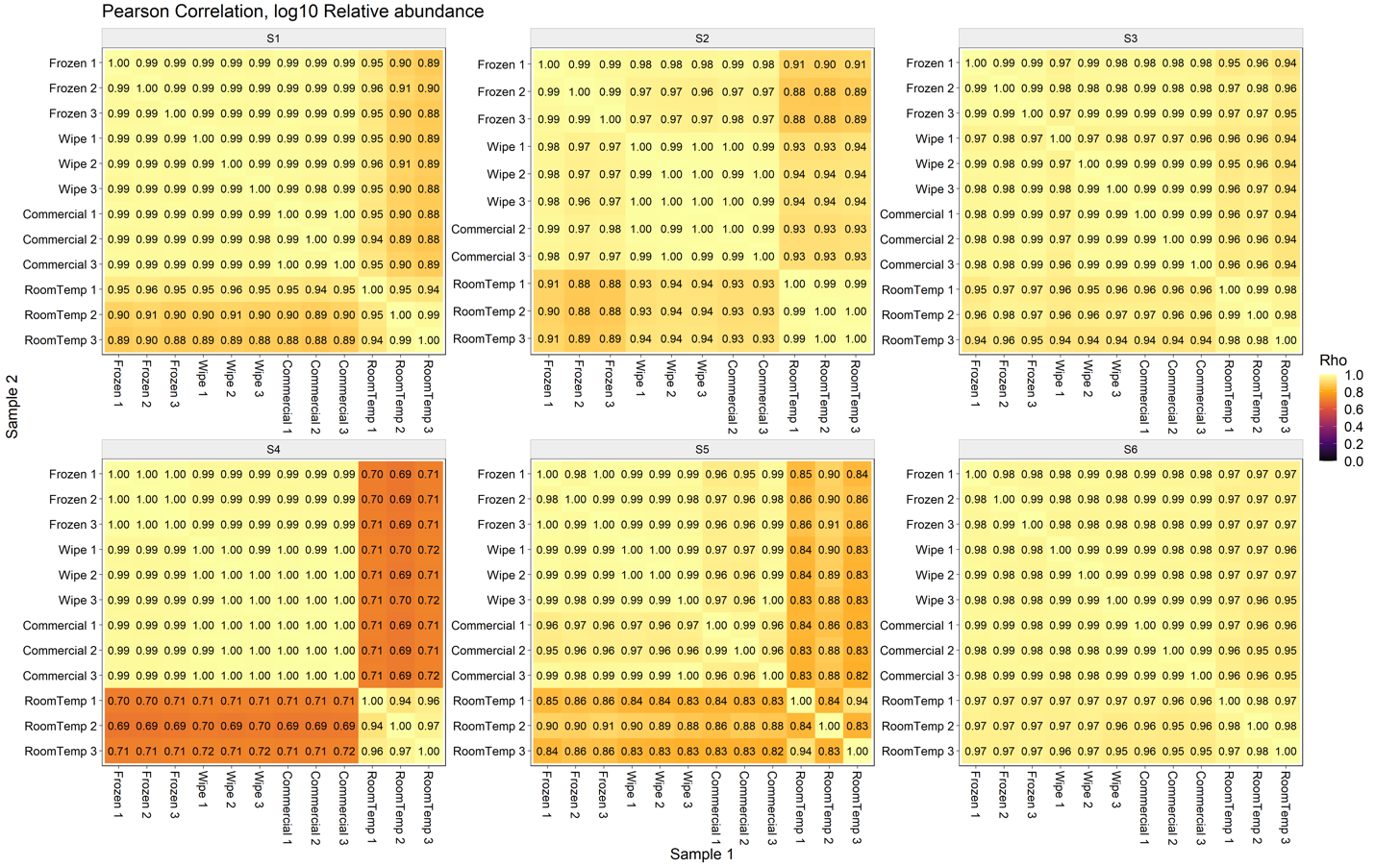

**Supplemental Figure 2. Taxonomic Pearson Correlation.** Pearson correlation of taxonomic relative abundances by sample type and subject.

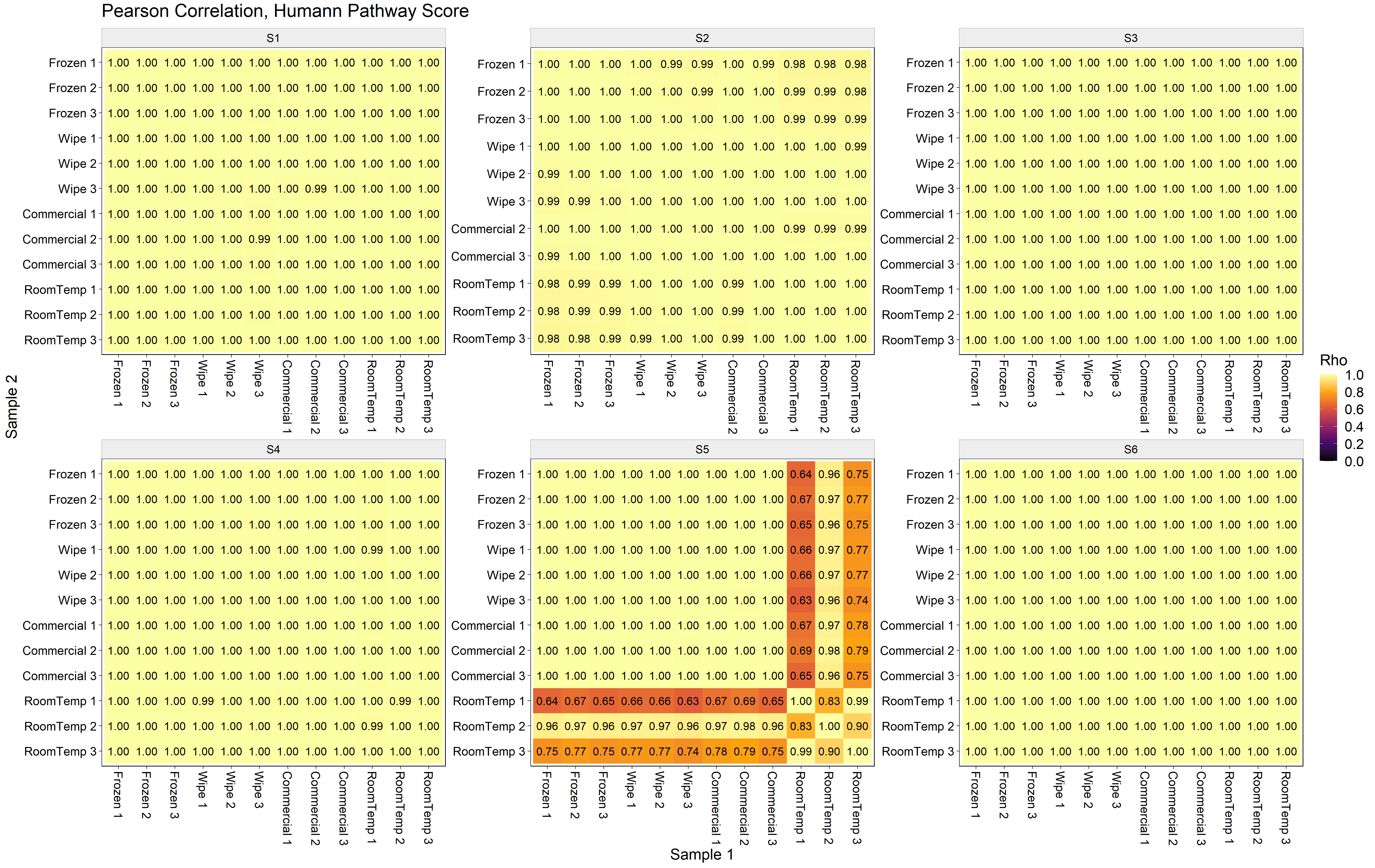

**Supplemental Figure 3. Functional Pathway Pearson Correlation.** Pearson correlation of HUMAnN pathway scores by sample type and subject.
